# Transcription promotes the interaction of the FAcilitates Chromatin Transactions (FACT) complex with nucleosomes in *S. cerevisiae*

**DOI:** 10.1101/376129

**Authors:** Benjamin J.E. Martin, Adam T. Chruscicki, LeAnn J. Howe

## Abstract

The FACT (FAcilitates Chromatin Transactions) complex is enriched on highly expressed genes, where it facilitates transcription while maintaining chromatin structure. How it is targeted to these regions is unknown. In vitro, FACT binds destabilized nucleosomes, supporting the hypothesis that FACT is targeted to transcribed chromatin through recognition of RNA polymerase-disrupted nucleosomes. In this study, we used high resolution analysis of FACT occupancy in *S. cerevisiae* to test this hypothesis. We demonstrate that FACT interacts with unstable nucleosomes *in vivo* and its interaction with chromatin is dependent on transcription by any of the three RNA polymerases. Deep sequencing of micrococcal nuclease-resistant fragments shows that FACT-bound nucleosomes exhibit differences in micrococcal nuclease sensitivity compared to bulk chromatin, consistent with a modified nucleosome structure being the preferred ligand for this complex. While the presence of altered nucleosomes associated with FACT can also be explained by the known ability of this complex to modulate nucleosome structure, transcription inhibition alleviates this effect indicating that it is not due to FACT interaction alone. Collectively these results suggest that FACT is targeted to transcribed genes through preferential interaction with RNA polymerase disrupted nucleosomes.

## INTRODUCTION

FACT (FAcilitates Chromatin Transactions) is an abundant and conserved complex that promotes DNA-dependent processes, such as transcription and DNA replication. In animals and plants, FACT is comprised of the subunits, Spt16 and SSRP1, while in *S. cerevisiae*, the role of SSRP1 is performed by two proteins, Pob3 and Nhp6 (Orphanides *et al*. 1999; Brewster *et al*. 2001; Formosa *et al*. 2001). FACT was originally identified through its ability to promote transcription elongation on a chromatin template in vitro (Orphanides *et al*. 1998). It was later shown to enhance TBP binding at nucleosomal sites and promote histone displacement from the promoters of inducible genes (Mason and Struhl 2003; Biswas *et al*. 2005). Consistent with these observations, yeast FACT (yFACT) destabilizes nucleosomes in vitro (Formosa *et al*. 2001; Rhoades *et al*. 2004; Xin *et al*. 2009; Valieva *et al*. 2016). Contrary to these data however, FACT has also been implicated in the stabilization of chromatin. In yeast, mutation of FACT subunits results in transcription initiation from cryptic sites within the bodies of genes, enhanced histone turnover, and a failure to re-establish chromatin following transcriptional repression (Jamai *et al*. 2009; Formosa 2012; Hainer *et al*. 2012; Voth *et al*. 2014). The seemingly opposing functions of FACT in both disrupting and stabilizing chromatin have been reconciled in a model in which FACT binds nucleosomes and maintains an altered nucleosome structure that allows RNA polymerase II (RNAPII) passage without histone loss (Formosa 2012).

FACT shows increased occupancy on highly expressed genes (Mayer *et al*. 2010; Feng *et al*. 2016; Pathak *et al*. 2018), and is targeted with RNAPII upon gene induction in both flies and budding yeast (Mason and Struhl 2003; Saunders *et al*. 2003; Duina *et al*. 2007; Nguyen *et al*. 2013; Vinayachandran *et al*. 2018). An unresolved question however, is how FACT is targeted to these regions. FACT interacts with multiple components of the transcription machinery, including RNA polymerases, the Paf1 complex, the chromatin remodeler, Chd1, and Cet1, a subunit of the capping enzyme, but whether any of these factors are required for global FACT occupancy has not been tested (Krogan *et al*. 2002; Squazzo *et al*. 2002; Simic *et al*. 2003; Tardiff *et al*. 2007; Sen *et al*. 2017). Moreover, various lines of evidence suggest that the recruitment of FACT may not be through direct interaction with the RNAPII transcriptional machinery. First, the timing of RNAPII and FACT recruitment to induced genes during heat shock differs (Vinayachandran *et al*. 2018). Second, FACT is also found at regions transcribed by RNAPI and III (Birch *et al*. 2009; Tessarz *et al*. 2014). Finally, histone mutants cause delocalization of FACT independently of RNAPII and other elongation factors (Duina *et al*. 2007; Lloyd *et al*. 2009; Nguyen *et al*. 2013; Pathak *et al*. 2018). Together, these data suggest that FACT is directed to transcribed regions through an indirect mechanism.

Several studies have demonstrated the interaction of FACT with nucleosomes in vitro, but only in the presence of a destabilizing stress. First, the HMG box protein, Nhp6, promotes the interaction of both human and yeast FACT with nucleosomes (Formosa *et al*. 2001; Ruone *et al*. 2003; Rhoades *et al*. 2004; Zheng *et al*. 2014; Valieva *et al*. 2016; Mccullough *et al*. 2018). Like other HMG box proteins, Nhp6 weakens histone-DNA contacts, potentially “priming” nucleosomes for FACT binding (Travers 2003; Hepp *et al*. 2017). Second, the introduction of a DNA double strand break promotes FACT binding to nucleosomes in vitro (Tsunaka *et al*. 2016). Third, curaxins, a class of drugs that promote unwrapping of nucleosomal DNA in vitro, “trap” FACT on chromatin (Gasparian *et al*. 2011; Safina *et al*. 2017; Nesher *et al*. 2018). Fourth, FACT binds hexasomes, made up of DNA, a H3-H3 tetramer and a single H2A-H2B dimer, but not nucleosomes (Wang *et al*. 2018). Finally, FACT is enriched in regions experiencing torsional stress (Safina *et al*. 2017), a force disruptive to nucleosomes (Teves and Henikoff 2014). These data, along with the disruptive effect of transcription on nucleosome structure (Kireeva *et al*. 2002; Schwabish and Struhl 2004; Kulaeva *et al*. 2010; Sheinin *et al*. 2013; Chang *et al*. 2014), support the intriguing hypothesis that FACT is targeted through preferential interaction with RNAP-destabilized nucleosomes. In this study, we tested this hypothesis using high resolution analysis of FACT occupancy in yeast. We demonstrate that the interaction of FACT with chromatin is dependent on transcription by any of the three RNA polymerases. Further, we show that FACT binds nucleosomes in vivo, preferentially interacting with genes with high nucleosome turnover. FACT-associated nucleosomes show altered nuclease sensitivity compared to bulk chromatin, which is consistent with FACT’s ability to modulate nucleosome structure, however the enhanced nucleus sensitivity is partially independent of transcription, suggesting that it is not due to FACT binding per se. Instead, these data support the model that FACT is targeted to transcribed regions through preferential interaction with RNA polymerase-destabilized nucleosomes.

## RESULTS

### Transcription promotes the interaction of FACT with chromatin

To identify pathways for targeting FACT to transcribed regions, we first sought to confirm co-localization of FACT with RNAPII. To this end, we performed ChIP-seq of HA-tagged Spt16 and the RNAPII subunit, Rpb3, from the same whole cell extracts with two independent biological replicates. As FACT undergoes a dramatic redistribution during S-phase (Foltman *et al*. 2013), we synchronized cells in G1 with α-factor. Our Spt16 ChIP-seq data agreed well with previously published FACT ChIP-seq experiments (Figure S1), and we observed a specific and striking co-localization of Spt16, but not an untagged control, with Rpb3 at individual loci (Figure 1A) and genome-wide (Figure 1B). In agreement with previous reports (Mason and Struhl 2003; Mayer *et al*. 2010; Liu *et al*. 2017; Pathak *et al*. 2018), we found FACT enriched over the bodies of transcribed genes (Figure 1C), with binding primarily occurring downstream of the +1 nucleosome.

**Figure 1:**
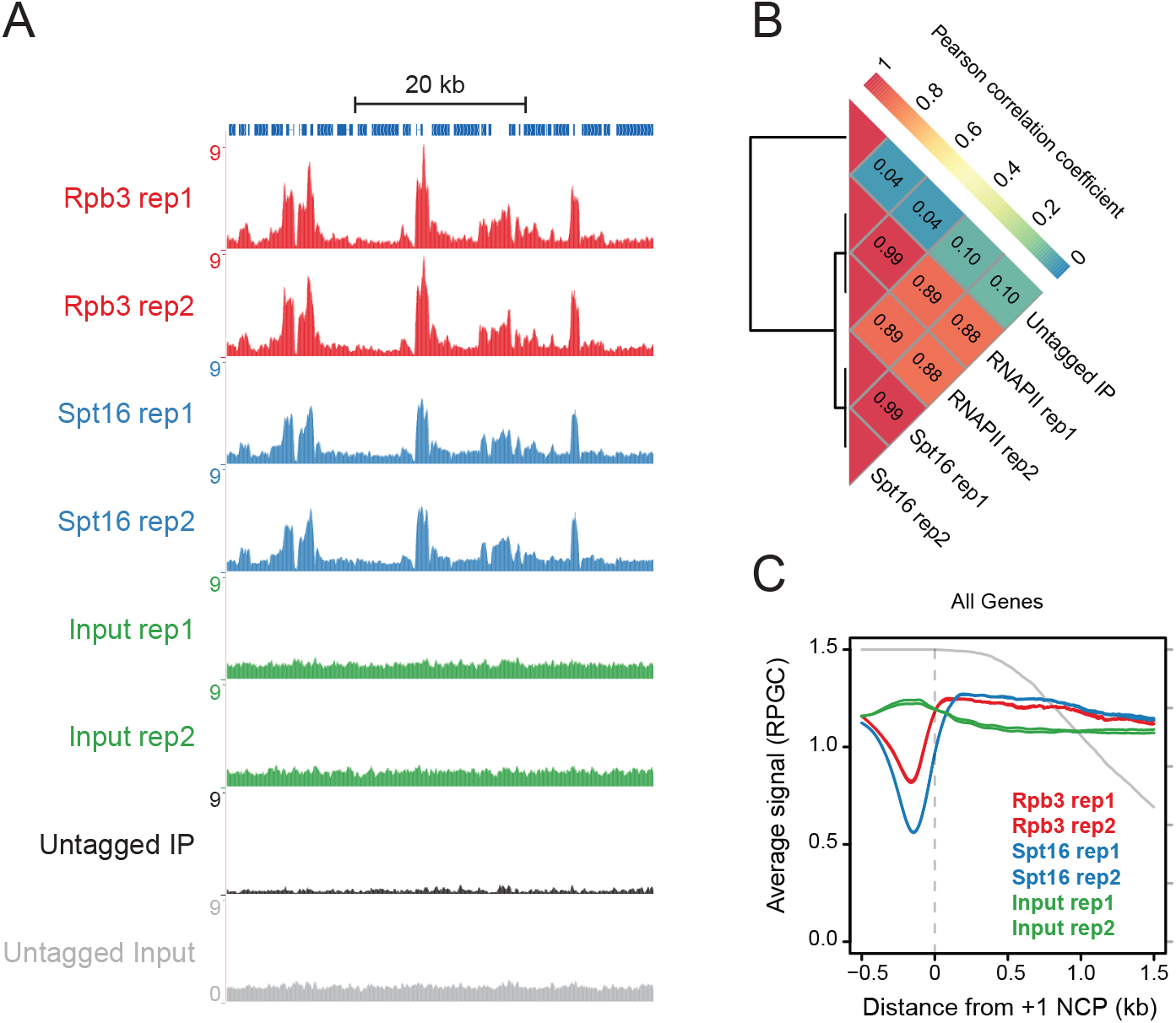
FACT and RNAPII co-localize. A) Genome browser tracks of sequence coverage (RPGC) from Rpb3 and Spt16 ChIP from sonicated chromatin (Input) and an untagged control over chrIV 300,000 - 350,000. The untagged IP was scaled by using spike-in exogenous DNA (see methods). B) Pearson correlation matrix for sequence coverage of Spt16, Rpb3 and untagged IPs across genome-wide 250bp bins. C) The average Rpb3, Spt16, and sonicated input sequence coverage (RPGC) relative to the +1 NCP of 5521 genes.

To determine whether transcription promotes FACT binding genome-wide, we first analyzed previously published data mapping RNAPII and Spt16 following depletion of Kin28, a kinase that facilitates promoter escape, using the rapamycin mediated “anchor away” technique (Wong *et al*. 2014). It should be noted that this analysis was conducted in cells grown in minimal media, which results in the previously observed, albeit unexplained, shift of RNAPII occupancy from the TSS to mid-gene when compared to cells grown in rich media (Warfield *et al*. 2017). Kin28 depletion results in a 5’ shift of both Rpb3 and Spt16, as well as loss of both proteins over gene bodies (Figures 2A and S2), consistent with targeting of FACT by transcription. This shift of FACT localization was not commented on previously, perhaps because the authors focused on a small subset of genes (Wong *et al*. 2014), but the 5’ shift of FACT upon Kin28 depletion appears as a general effect across the 5521 genes analyzed here.

**Figure 2:**
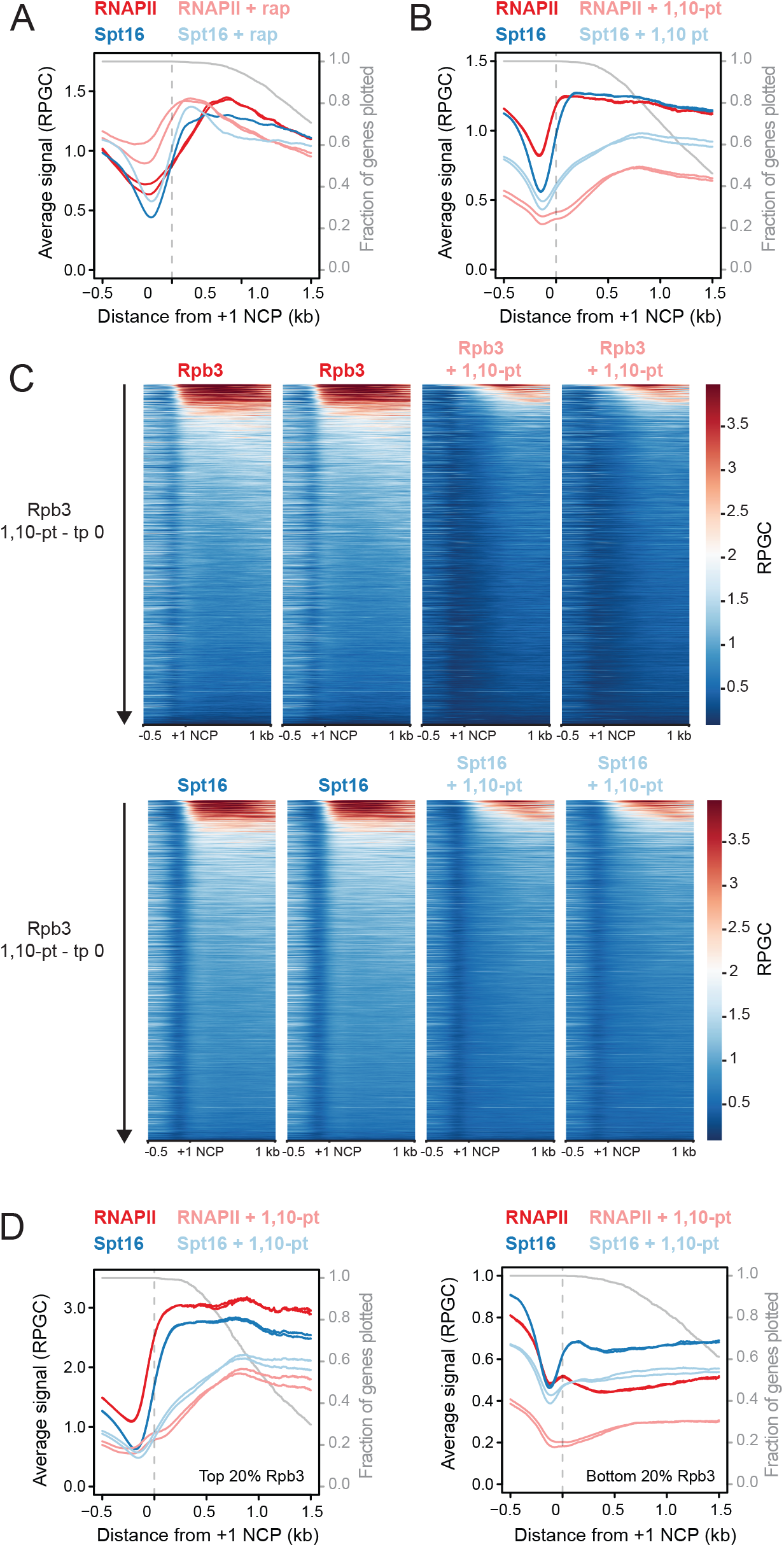
The interaction of FACT with chromatin is dependent on transcription. A) Rpb3 and Spt16 sequence coverage (RPGC), from cells expressing Kin28 with an FRB tag that was “anchored away” in the presence of rapamycin (rap), relative to the +1 NCP of 5521 genes. B) As in (A) except from cells with and without treatment with 1,10-phenanthroline (1,10-pt). Fragment coverage was normalized using “spiked-in” control DNA. C) Rpb3 and Spt16 sequence coverage (RPGC) relative to 5521 +1 NCPs (Chereji et al. 2018) prior to or following 15 minute treatment with 1,10-pt, represented by heatmap. D) As in (B) but for the top and bottom 20% of transcribed genes (1105 genes), as determined by Rpb3 binding.

To more directly test whether FACT requires RNAPII for binding to transcribed genes we next treated cells with the general transcription inhibitor 1,10-phenanthroline monohydrate (1,10-pt) (Grigull *et al*. 2004) and analyzed Spt16 and RNAPII occupancy by ChIP-seq. Importantly, “spike-in” DNA controls were added immediately following DNA elution to normalize the number of DNA fragments recovered from treated to untreated cells (Chen *et al*. 2015). Although the mechanism of action of 1,10-pt is not fully understood, short treatment (15 minutes) resulted in depletion of RNAPII across gene bodies, with the greatest effect over the 5’ regions (Figures 2B and C). Similar to RNAPII, treatment of cells with 1,10-pt also depleted Spt16 from gene bodies and shifted Spt16 occupancy from 5’ to 3’ genic regions at both highly and poorly transcribed genes (Figures 2B-D), indicating that the interaction of FACT with DNA requires transcription. Supporting a strong dependence of FACT targeting on RNAPII, the changes in Spt16 and Rpb3 occupancy upon transcription inhibition were highly correlated genome-wide (Figure S3). Collectively, these results demonstrate that FACT targeting to transcribed regions occurs as a consequence of RNA polymerase activity.

While the above data is consistent with a model in which FACT is targeted through physical interaction with the RNAPII transcriptional machinery, several contradictions with this model exist. First, although FACT and RNAPII levels tightly correlated across gene bodies, poorly expressed genes exhibited higher binding of Spt16 relative to RNAPII than highly expressed genes (Figure 2D). This observation is better visualized in Figure S4A, which compares the ratio of Spt16 to Rpb3 with Rpb3 levels. Such enrichment at poorly expressed genes was not observed for Spt5, an elongation factor that interacts directly with RNAPII (Figure S4B). Second, following transcription inhibition, the magnitude of Spt16 depletion was less than for Rpb3, even at 5’ genic regions where Rpb3 depletion is most evident (Figure 2B), suggesting that FACT remains transiently associated with chromatin following RNAPII passage. Third, the length of DNA fragments co-precipitating with Spt16 and Rpb3 differed markedly (Figure S5A), with Rpb3-associated DNA being smaller than that associated with Spt16. Notably, the differences in fragment sizes were observed at both lowly and highly transcribed regions of the genome and across gene bodies (Figure S5B and C), confirming that the differences in fragment size are occurring at regions of co-localization, at least as observed across a population of cells. The differences in fragment sizes are unlikely due to technical variation in sample treatment as these ChIPs were performed from the same cell extracts, the DNA purifications and sequencing libraries were prepared in parallel, and the libraries were indexed, pooled, and sequenced together. Instead, the differences in DNA fragment size likely indicate that the majority of Rpb3 and Spt16 do not purify as a single nucleoprotein complex. Finally, transcription inhibition also resulted in depletion of FACT from RNAPI and III-transcribed regions (Figures S6, S7 and S8). Collectively, these data support a model whereby transcription promotes the interaction of FACT with chromatin via an indirect mechanism.

### FACT binds polymerase-destabilized nucleosomes in vivo

FACT binds disrupted nucleosomes in vitro (Formosa *et al*. 2001; Ruone *et al*. 2003; Zheng *et al*. 2014; Tsunaka *et al*. 2016; Mccullough *et al*. 2018; Nesher *et al*. 2018; Wang *et al*. 2018), supporting the hypothesis that it is targeted to transcribed genes through interaction with RNA polymerase-destabilized nucleosomes. To confirm that FACT binds nucleosomes in vivo, we purified FACT via a TAP tag on Pob3. Figure 3A shows that FACT co-purified with stoichiometric levels of histones H2A, H2B, H3 and H4. Immunoprecipitation of H2B from purified FACT co-precipitated all four core histones (Figure 3B), indicating that FACT interacts with intact histone octamers as opposed to select histone pairs. To test whether these histones were nucleosomal, we ChIPped HA-tagged Spt16 from MNase-treated extracts prepared from formaldehyde cross-linked cells. The co-purifying DNA was subjected to paired-end sequencing and a heatmap showing the location of sequence reads across genes sorted by RNAPII occupancy showed that the positioning of FACT-associated DNA closely matched that of input nucleosomes (Figure 3C). This is more evident in Figure S9, in which the total number of sequence fragments recovered were normalized between genes to compensate for increased Spt16 occupancy at highly expressed genes. Despite the similarities between input and FACT-bound chromatin, noticeable differences were observed including increased recovery of sequence fragments from highly expressed genes, depletion and shift of the +1 nucleosome peak, and enrichment of FACT over gene bodies (Figure 3D). Thus, despite the reported inability of FACT to bind nucleosomes in the absence of destabilizing stress in vitro (Tsunaka *et al*. 2009; Wang *et al*. 2018), our results suggest that FACT can bind nucleosomes on transcribed genes in vivo.

**Figure 3:**
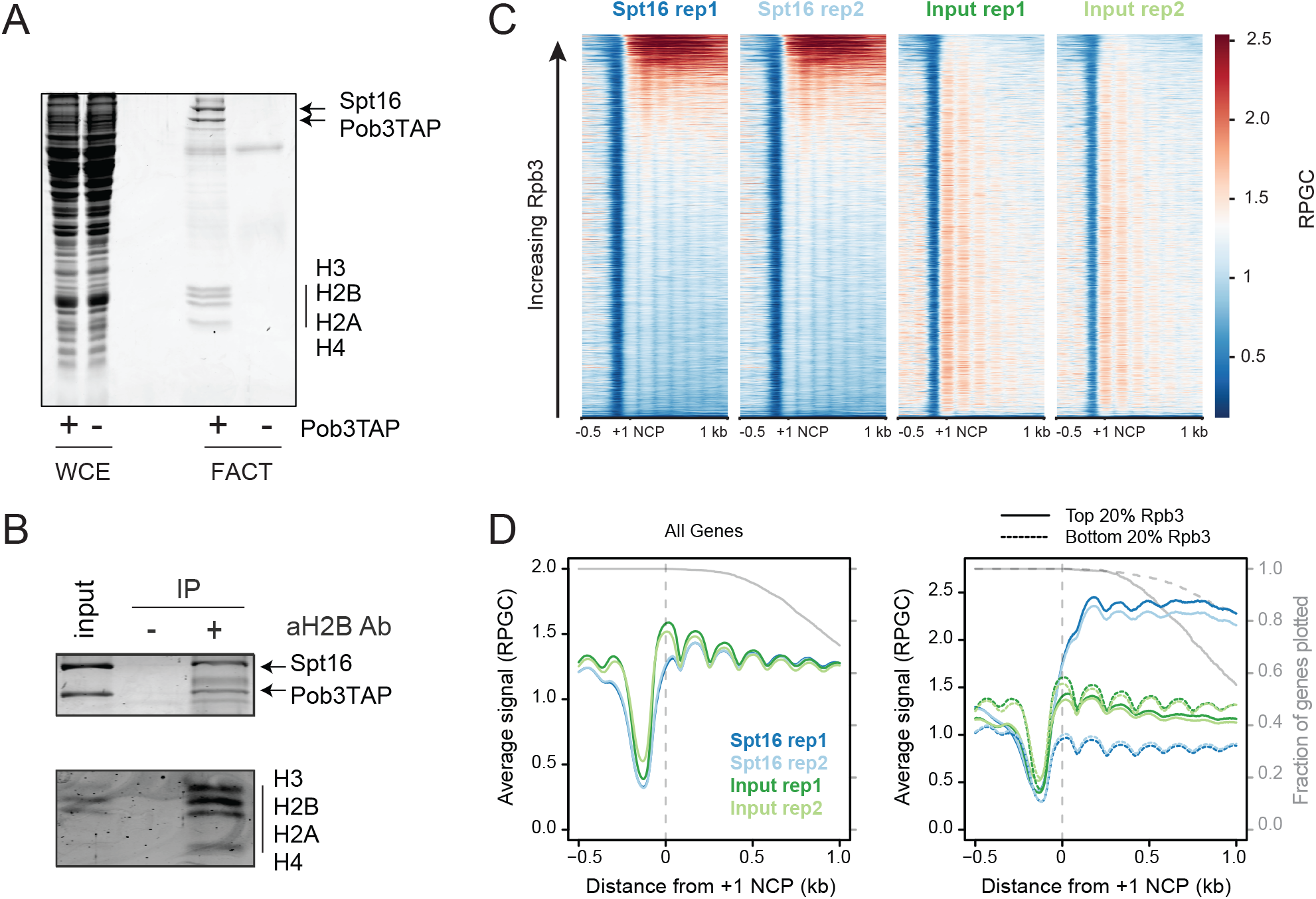
FACT binds nucleosomes in vivo. A) FACT, purified from a strain expressing TAP-tagged Pob3 (+), was subjected to SDS PAGE analysis with coomassie staining. And untagged strain (−) was used as a negative control. B) FACT and associated histones were purified from whole cell extracts via a TAP tag on Pob3. Purified FACT was eluted, by cleavage of the TAP tag with TEV protease, and subjected to immunoprecipitation with antibodies specific for H2B. The proteins in input and immunoprecipitated fractions were visualized by coomassie staining of an SDS PAGE gel. C) Spt16 and MNase input sequence coverage (RPGC) relative to 5521 +1 NCPs (Chereji et al. 2018) represented by heatmap. D) As in (C), but data represented as the average sequence coverage (RPGC) overlapping 5521 +1 NCPs and the top and bottom 20% of Rpb3 bound genes (1105 genes each).

We next asked whether FACT preferentially interacts with unstable nucleosomes. One hallmark of unstable nucleosomes is increased histone turnover, and indeed, Spt16 occupancy correlated with replication independent (RI) histone turnover (Figure 4A, Spearman rank correlation coefficient of 0.47) (Dion *et al*. 2007). Histone loss is also associated with high levels of transcription, so to test if FACT correlated with histone turnover independently of RNAPII-levels, we employed a technique called partial correlation, a statistical method that calculates the association between two factors after eliminating the effects of other variables (Kim 2015). Importantly, this technique allows estimation of the direct interaction between two factors while controlling for a confounding variable. Partial correlation analysis revealed that RI histone turnover over gene bodies directly correlated with Spt16 and not Rpb3 (Figure 4A), demonstrating that FACT co-localizes with RI histone turnover independently of transcription. To assess this relationship another way, we ordered genes by Rpb3 occupancy, and in a sliding window of 500 genes assessed the mean level of RI histone turnover for the 100 genes with the highest or lowest Spt16 occupancy relative to Rpb3 (Figure 4B). Aside from very highly transcribed genes, we found RI histone turnover increased at genes with higher Spt16 relative to Rpb3 occupancy (Figure 4C). Although we cannot rule out the possibility that the link between Spt16 occupancy and histone turnover is due to FACT promoting RI histone turnover, the demonstrated ability of FACT to suppress histone turnover *in vivo* argues that this is not the case (Jamai *et al*. 2009). Indeed, analyzing published MNase-seq data in *spt16* mutant cells (True *et al*. 2016) revealed that FACT inactivation leads to preferential loss of nucleosomes at these genes (Figure 4D), supporting a model whereby FACT preferentially binds to regions of destabilized nucleosomes where it acts to stabilize chromatin.

**Figure 4:**
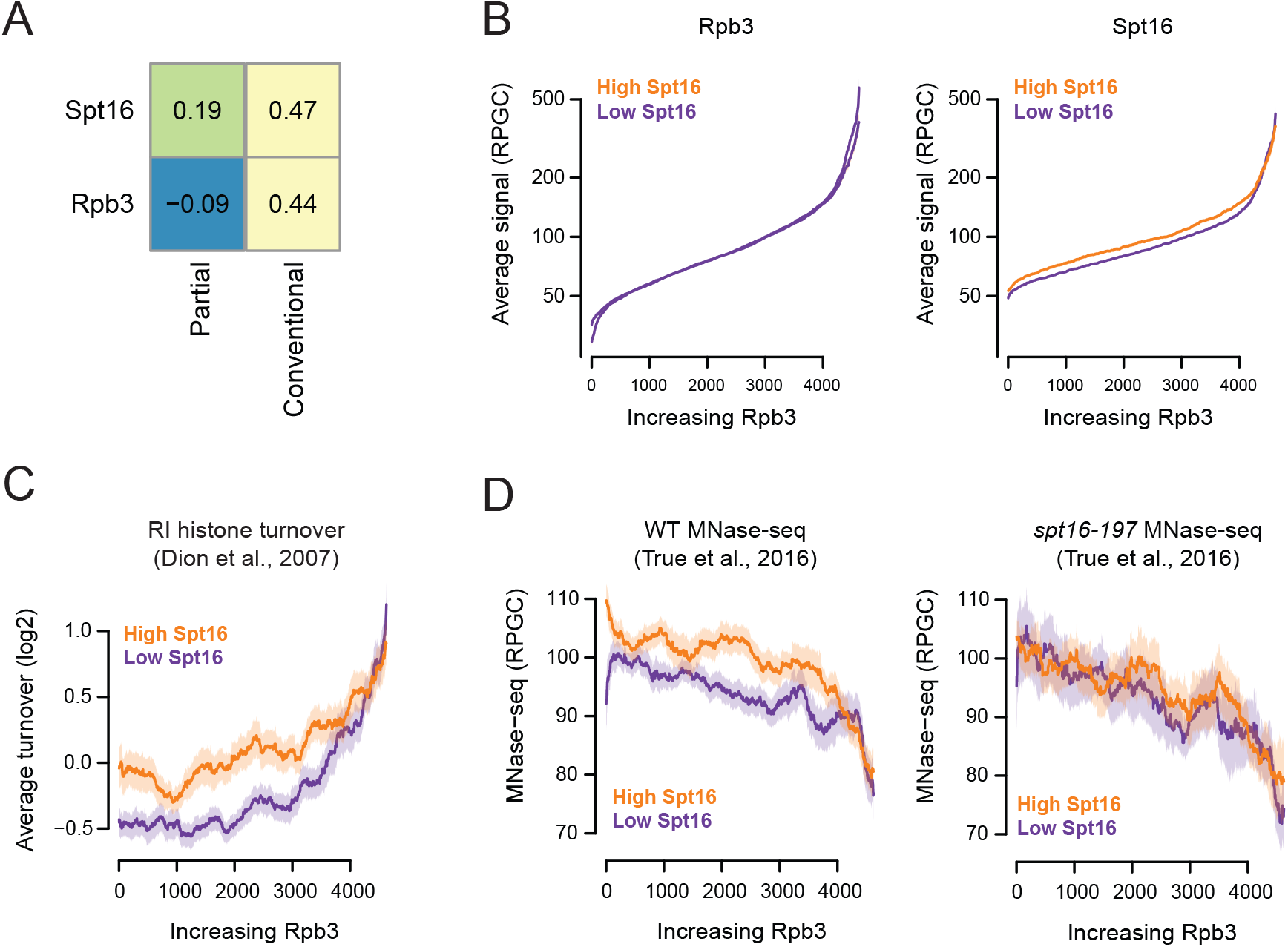
FACT co-localizes with RI histone turnover independently of RNAPII. A) Gene-based conventional and partial Spearman correlation coefficients represented by heatmap. B) Genes, as in (A), were ordered by Rpb3 levels and in 500 gene sliding windows the top and bottom 20% of Spt16-Rpb3 were designated as high and low Spt16 groups. Mean enrichment of Rpb3, Spt16 are plotted with the shaded region representing the 95% confidence interval. C,D) Using the same gene windows as in (B), RI histone turnover (Dion et al., 2007) (C) and MNase-seq coverage from wildtype and spt16-197 strains following 45 minute shift to the restrictive temperature (D) are plotted with the shaded region representing the 95% confidence interval.

The use of paired-end sequencing in our analyses afforded us the opportunity to examine the nuclease sensitivity, and thus potential alterations, of FACT-bound nucleosomes. While FACT-associated DNA exhibited resistance to MNase, consistent with the binding of nucleosomes, specific alterations from bulk chromatin were observed (Figure 5A). FACT-bound DNA fragments, but not those from an untagged control, exhibited increased levels of longer (200 - 260 bp) and shorter (100 - 130 bp) than mononucleosome sized (140 - 160 bp) fragments. These differences were unlikely to have arisen from technical variation in sample processing as the ChIPs and inputs were processed in parallel, indexed, and pooled together for sequencing. To determine where the altered, FACT-bound nucleosomes were located, we used two-dimensional occupancy plots to visualize the boundaries of MNase-resistant fragments associated with FACT or with bulk chromatin, relative to the +1 nucleosomes of RNAPII transcribed genes. Shown in Figure 5C, these plots simultaneously display DNA sequencing data as: 1) a heatmap of the total counts of the left (left panels) and right (right panels) boundaries of MNase protection (as designated in Figure 5B), 2) the total length of the MNase-resistant fragments (Y-axis) and 3) position of the boundary relative to the dyad of the +1 nucleosomes of 5521 genes (X-axis). Analysis of bulk chromatin showed peaks of left boundary signal at ~−75 bp and right boundary signal at ~+75 bp relative to +1 nucleosomes on the X-axis and ~150 bp on the Y-axis, consistent with well positioned, +1 mononucleosomes protecting ~150 bp of DNA (Figure 5C, top panels). Also evident, albeit fainter, was a ~−75 bp left boundary signal and a ~+250 bp right boundary signal at ~315 bp on the Y-axis, indicative of di-nucleosomes containing both the +1 and +2 nucleosomes. Of additional note, signals were also observed below 150 bp on the Y-axis, that slope toward the nucleosome dyads. MNase preferentially digests linker DNA, but transient unwrapping of DNA from the octamer surface can result in “chewing” of the DNA ends. Thus, these lower signals likely represent MNase trimming of nucleosomes from either direction.

**Figure 5:**
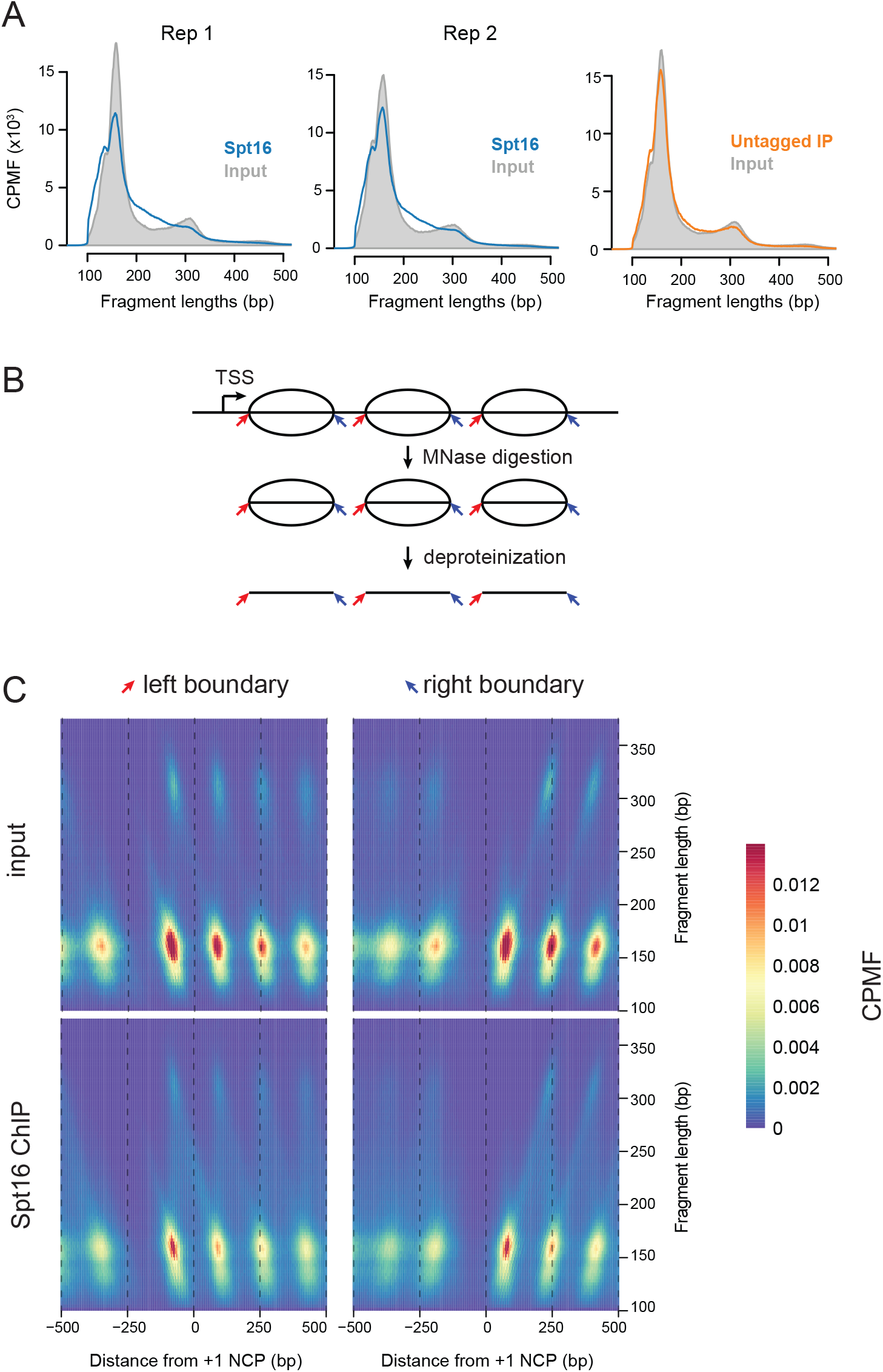
FACT-bound nucleosomes exhibit altered MNase sensitivity. A) Histograms depicting DNA fragment lengths recovered from input, Spt16-6HA, and untagged control ChIPs from MNase-digested chromatin B) Designation of left (red arrows) and right (blue arrows) nucleosome fragment boundaries identified through MNase digestion of chromatin. C) Two dimensional plots of positions of nucleosome boundaries in input and Spt16 ChIP from MNase digested chromatin, relative to the +1 nucleosome of 5521 annotated genes in S. cerevisiae. the sum of fragment boundary counts are indicated as a heatmap, the sequence fragment lengths (split into 3 bp bins) is plotted ont he Y-axis and the position of nucleosome boundaries relative to the +1 nucleosome is plotted on the X-axis.

Performing 2D analysis of DNA fragment boundaries from FACT-associated nucleosomes revealed many differences from bulk nucleosomes (Figure 5C, lower panels). First, nucleosomes bound to FACT showed an increased proportion of “trimmed” nucleosomes. These smaller fragments could also represent hexasomes, reported to protect ~110 bp from MNase (Arimura *et al*. 2012), as FACT has been shown to form a ternary complex with a single dimer of histones H2A-H2B and a H3-H4 tetramer in vitro (Wang *et al*. 2018). Interestingly, equivalent MNase trimming of Spt16-bound fragments was observed on the promoter-proximal (left boundary) and promoter-distal (right boundary) edges of nucleosomes, suggesting that the direction of transcription had little impact on the presence of these “sub-nucleosomes”.

Another interesting result from our 2D analysis is that discrete signals at 315 bp, representative of di-nucleosomes, were less evident in FACT-associated DNA. Instead, a continuum of fragment sizes from mono to di-nucleosomes was observed, suggesting that FACT can alter typical spacing between nucleosomes. Indeed, DNA fragments in the 200 to 260 bp range could be indicative of “overlapping di-nucleosomes”. Such structures, composed of a hexasome and a nucleosome, have been demonstrated in vitro by many labs (Ulyanova and Schnitzler 2005; Engeholm *et al*. 2009; Kato *et al*. 2017), and indeed, the susceptibility of FACT-associated nucleosomes to lose an H2A/H2B dimer (Belotserkovskaya *et al*. 2003) may make these nucleosomes more prone to forming such structures. Consistent with this, Figure 5A shows that FACT-bound nucleosomes were depleted in the canonical di-nucleosome signal observed in total chromatin.

While, the altered nuclease sensitivity of Spt16-bound nucleosomes is consistent with a preference of FACT for transcription-disrupted nucleosomes, FACT is also reported to alter nucleosome structure *in vitro* (McCullough et al., 2018; Xin et al., 2009), and thus a modified nucleosome structure could also be a consequence of FACT binding. To differentiate between these two possibilities, we rationalized that, if FACT modifies nucleosome structure on its own, then the altered pattern of digestion should be independent of the destabilizing stress of transcription. To test this, we divided input and Spt16 ChIP DNA fragments into three bins [100-130 bp (purple lines), 140-160 bp (teal lines) and 200-260 bp (orange lines)] and plotted the abundance of each relative to the +1 nucleosomes of highly expressed genes (dark lines) and poorly expressed genes (light lines) (Figure 6). This analysis revealed Spt16 co-precipitated increased amounts of shorter (100 – 130 bp) and longer (200 – 260 bp) sized DNA fragments from highly expressed genes compared to poorly expressed genes. Interestingly, the major species of larger FACT-bound fragments predominantly occurred upstream and overlapping the +2 NCP, suggesting that the +1 NCP has shifted downstream to create an overlapping di-nucleosome of the +1 and +2 NCPs. The increased levels of alternate fragments at highly expressed genes was diminished upon transcription inhibition. Collectively these results indicate that altered MNase sensitivity of FACT-bound nucleosomes is dependent on transcription, suggesting that nucleosome disruption is primarily a cause as opposed to consequence of FACT binding. These data are consistent with a model in which FACT is targeted to active genes through preferential interaction with RNAP-disrupted nucleosomes.

**Figure 6:**
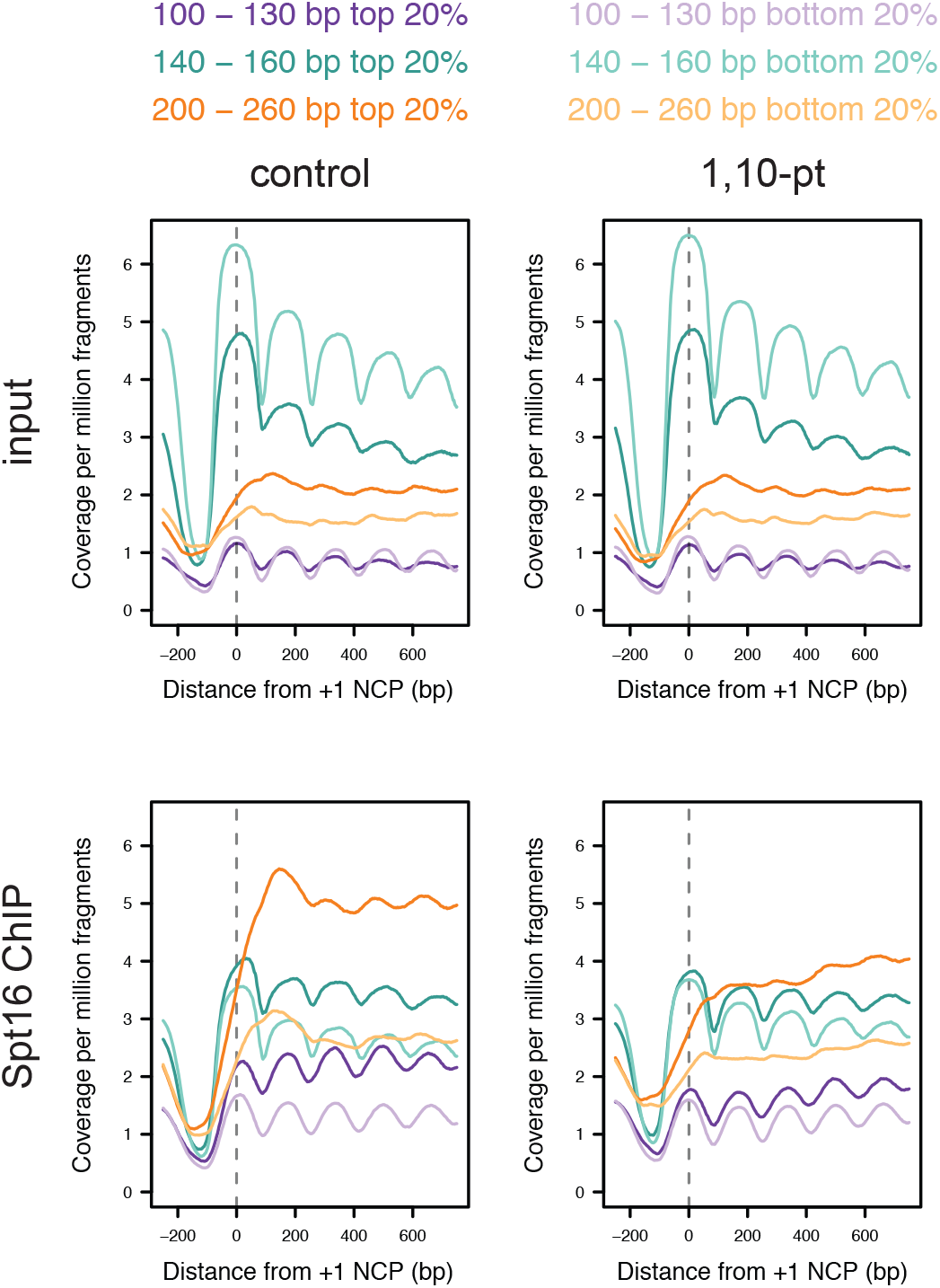
Transcription enhances nuclease senstivity of FACT-bound nucleosomes. MNase input and Spt16 ChIP-seq sequence for cells with and without treatment with 1,10-pt, relative to the +1 NCP of the top and bottom 20% of transcribed genes (1105 genes each). Sequence fragments were selected and coverage per million fragments (of all sizes) is plotted.

## DISCUSSION

In this study, we show that the FACT complex binds nucleosomes in vivo and is recruited to chromatin as a consequence of transcription by any of the three RNA polymerases. These data, together with extensive evidence that FACT binds destabilized nucleosomes in vitro, are consistent with a model in which FACT is targeted to transcribed regions through preferential interaction with RNAP-destabilized nucleosomes. Individual domains of FACT subunits have been shown to bind DNA and all four core histones in vitro (Stuwe *et al*. 2008; Vandemark *et al*. 2008; Winkler *et al*. 2011; Hondele *et al*. 2013; Kemble *et al*. 2013; Hoffmann and Neumann 2015; Kemble *et al*. 2015; Tsunaka *et al*. 2016). The unwrapping of nucleosomal DNA by RNA polymerases could reveal high affinity binding sites that stabilize the binding of FACT at transcribed genes.

Our results show that Spt16-associated nucleosomes exhibit altered sensitivity to MNase, which is indicative to FACT binding to atypical nucleosome structures, including potentially overlapping di-nucleosomes. While the function of overlapping di-nucleosomes has not been investigated (Lowary and Widom 1998; Ulyanova and Schnitzler 2005; Engeholm *et al*. 2009; Kato *et al*. 2017), one possibility may be to increase nucleosome mobility on transcribed genes. Nucleosomes restrict the movement of their neighbors (Kornberg and Stryer 1988; Zhang *et al*. 2011), but facilitating formation of overlapping nucleosomes would increase flexibility of movement without the need for octamer eviction, which coincides with the ability of FACT to both promote transcription while stabilizing chromatin structure. The altered sensitivity of FACT-associated nucleosomes is largely dependent on active transcription, which is consistent with it being a cause, as opposed to a consequence of FACT binding. However, it should be noted that multiple studies have shown that FACT can disrupt nucleosome structure in vitro (Xin *et al*. 2009; Valieva *et al*. 2016), and thus we cannot rule out the possibility that FACT further alters chromatin structure once bound.

In addition to functioning in transcription, FACT plays an essential role in DNA replication. During S phase, FACT relocates to newly replicated chromatin (Foltman *et al*. 2013; Alabert *et al*. 2014), where it interacts with multiple components of the replication machinery (Wittmeyer and Formosa 1997; Gambus *et al*. 2006; Vandemark *et al*. 2006). Moreover, mutation of FACT results in sensitivity to hydroxyurea (HU), delayed S phase progression (Schlesinger and Formosa 2000; Herrera-moyano *et al*. 2014) and defects in chromatin assembly on nascent DNA (Yang *et al*. 2016). Finally, mutation of FACT results in dependence on the S phase checkpoint for viability (Schlesinger and Formosa 2000). However, despite extensive data supporting a requirement for FACT in DNA replication, its actual function in this process is unclear. Part of this mystery is routed in confusion over the molecular function of FACT. While FACT was originally proposed to be a histone chaperone that deposits free histones on DNA (Belotserkovskaya *et al*. 2003), data overwhelmingly suggests that FACT instead binds intact nucleosomes (Ruone *et al*. 2003; Tsunaka *et al*. 2009; Xin *et al*. 2009; Winkler *et al*. 2011; Tsunaka *et al*. 2016; Valieva *et al*. 2016; Valieva *et al*. 2017). Chromatin assembled on newly replicated DNA undergoes a maturation step shortly following DNA replication (Fennessy and Owen-Hughes 2016; Vasseur *et al*. 2016), raising the possibility that FACT recognizes newly formed, unstable nucleosomes on nascent DNA, and akin to its role in transcription, stabilizes these nucleosomes until maturity.

## MATERIALS AND METHODS

### Yeast strains and antibodies

All strains used in this study were isogenic to S288C. The Pob3TAP and Spt16HA_6_ were derived from FY602 (*MATa his3Δ200 lys2-128δ leu2Δ1 trp1Δ63 ura3-52*), a generous gift of Fred Winston. Yeast culture, genetic manipulations, and strain verification were performed using standard protocols. Antibodies used were from Roche [11 583 816 001 (HA)], Biolegend [665004 (Rpb3)], Active Motif [39237 (histone H2B)], and Millipore [PP64 (IgG)].

### Spt16 Chromatin ImmunoPrecipitation (ChIP) sequencing

Yeast cells were arrested in G1 by treatment with 10 μM alpha factor for three hours (synchronization confirmed by inspection of cell morphology under the microscope). Transcription inhibition was performed by treatment with 400 μg/mL 1,10-pt for 15 minutes. Crosslinking was performed with 1% (vol/vol) formaldehyde for 15 minutes at room temperature and quenched with 125 mM glycine for 15 minutes.

Sonicated ChIP-seq was performed essentially as described previously (Lawrence *et al*. 2017). Briefly, cells were disrupted by bead beating, chromatin was sonicated to an average DNA length of 250 bp (Biorupter, Diagenode), and parallel immunoprecipitations were performed using α-HA or α-Rpb3 antibodies.

MNase ChIP-seq was performed essentially as described previously (Maltby *et al*. 2012). Briefly, cells were disrupted by bead beating, chromatin was digested to mononucleosomes with MNase, and immunoprecipitations were performed using α-HA antibody. Sequencing libraries were prepared as described previously (Maltby *et al*. 2012; Martin *et al*. 2017). Exogenous DNA spike-ins were added to ChIP-seq eluates () in order for quantification of global changes in IP efficiency.

Sequencing libraries were prepared essentially as described previously (Maltby *et al*. 2012). Briefly, 2 ng of IP or input material was end-repaired, A-tailed, and adapters ligated, before PCR amplification with indexed primers. We performed 10 and 11 rounds of PCR amplification for MNase and sonication ChIP-seq experiments respectively. All DNA purification steps used SPRI beads. Samples were pooled, gel-purified (50 bp - 600 bp region) and sequenced on an Illumina HiSeq 2500.

### Data analysis

Sequenced reads were trimmed for adapter sequences using cutadapt (http://cutadapt.readthedocs.io/en/stable/). Trimmed reads were then aligned to the Saccer3 genome using bowtie2 (Langmead and Salzberg 2012), and filtered for paired reads with a mapping quality of at least 5 using SAMtools (Li *et al*. 2009)). Fragments mapping to synthetic spike in sequences were used to scale ChIP-seq datasets following 1,10-pt transcription inhibition. Reads Per Genome Coverage (RPGC) bigwig tracks were generated using deeptools (Ramirez *et al*. 2014; Ramirez *et al*. 2016). Size selected Counts Per Million Fragments (CPMF) tracks were generated using a custom bash script using AWK, the BEDtools (Quinlan and Hall 2010), and the UCSC bedgraphtobigwig function (Kent *et al*. 2010). Genome browser shots were generated using the UCSC genome browser (Kent *et al*. 2002).

Genome wide coverage in genome wide windows for ChIP-seq data was generated using deeptools, and heatmaps and scatter plots were plotted in R using pheatmap and smoothscatter functions respectively. Fragment length histograms were plotted in R, and fragments overlapping specific genomic regions were selected for using BEDtools. Coverage heatmaps were generated using deeptools. For average plots and row-normalized heatmaps, feature-aligned matrixes were constructed using deeptools and plots were then generated in R using base plots or pheatmap functions respectively. For average plots, to avoid effects due to varying gene lengths, plots only include data until the TTS, with the fraction of genes plotted at a given position indicated by a grey line. The MNase 2D heatmaps were generated using AWK and BEDtools, with the final heatmaps were plotted in R using pheatmap.

For gene body comparisons of Rpb3 and Spt16, gene bodies were defined as +73 bp downstream of the +1 dyad, to avoid initiation effects, to the TTS. To avoid gene-length effects, only genes (+1 dyad to TSS) longer than 500 bp were analyzed, but similar results were seen across all gene lengths. ChIP-seq fragments overlapping gene bodies were counted per million fragments and scatter plots generated using the smoothscatter function in R. Partial correlations were calculated using the ppcor function in R (Kim 2015). For histone turnover data (Dion *et al*. 2007), average values over gene bodies were calculated using the java genomics toolkit (https://github.com/timpalpant/java-genomics-toolkit). For MNase-seq data from Wildtype and *spt16-197* strains, DNA fragments (True *et al*. 2016) overlapping gene bodies were counted per million fragments. For gene windows, genes were ordered by Rpb3 levels in a 500-gene sliding window. For each window the top and bottom 100 genes for Spt16 minus Rpb3 were selected and the mean value plotted. The 95% confidence interval was bootstrapped and plotted as the shaded area in the graph. Colour schemes were generated using the RColorBrewer Package in R.

### Published datasets

For histone turnover data (Dion *et al*. 2007), probe intensity values were converted to Saccer3 using the UCSC liftover tool and conversion to wig format, linear interpolation to a maximum of 500 bp, and average values over gene bodies were calculated using the java genomics toolkit (https://github.com/timpalpant/java-genomics-toolkit). Wildtype and *spt16-197* MNase-seq datasets (True *et al*. 2016) were downloaded from SRA project SRP073244. Spt16 ChIP-seq datasets were downloaded from SRA projects SRP055441 (Feng *et al*. 2016), SRP018874 (Foltman *et al*. 2013), SRP073244 (True *et al*. 2016), SRP036647 (Wong *et al*. 2014). The Spt16 ChIP-exo dataset was downloaded from SRA project SRP106497 (Vinayachandran *et al*. 2018). Spt5 and Rpb1 ChIP-seq datasets (Baejen *et al*. 2017) were downloaded from SRP071780. Sequenced reads were processed as described above.

Transcription Start Site (TSS) annotations are from Park et al (PARK *et al*. 2014). Plus One Nucleosome and Transcription Termination Site (TTS) are from (Chereji *et al*. 2018). Yeast genome features were downloaded from http://www.yeastgenome.org/

### FACT purification

Pob3TAP and its associated proteins were purified from one litre of yeast cells as described (Lambert *et al*. 2009). For micrococcal nuclease digestion, magnetic beads containing purified FACT-nucleosome complexes were washed in 50 mM Tris pH 8, 4 mM MgCl_2_, 2 mM CaCl_2_ and digested with increasing amounts of nuclease. Reactions were stopped by washing the beads in 50 mM Tris pH 8, 4 mM MgCl_2_, 10 mM EDTA and the DNA was purified using phenol:chloroform:isoamyl alcohol extraction followed by ethanol precipitation.

## Data availability

Strains and plasmids are available upon request. The ChIP-seq data generated for this study have been deposited in the Gene Expression Omnibus (GEO) database under accession numbers GSE111426 and GSE110286. Supplementary figures have been uploaded to figshare.

## Acknowledgments

We acknowledge Tim Formosa for his constructive feedback. This work was supported by a Natural Sciences and Engineering Research Council (NSERC) Discovery Grant awarded to L.J.H. B.J.E.M was supported by graduate student fellowships from NSERC.

